# *WOLO* : Wilson Only Looks Once – Estimating ant body mass from reference-free images using deep convolutional neural networks

**DOI:** 10.1101/2024.05.15.594277

**Authors:** Fabian Plum, Lena Plum, Corvin Bischoff, David Labonte

## Abstract

Size estimation is a hard computer vision problem with widespread applications in quality control in manufacturing and processing plants, livestock management, and research on animal behaviour. Image-based size estimation is typically facilitated by either well-controlled imaging conditions, the provision of global cues, or both. Reference-free size estimation remains challenging, because objects of vastly different sizes can appear identical if they are of similar shape. Here, we explore the feasibility of implementing automated and reference-free body size estimation to facilitate large-scale experimental work in a key model species in sociobiology: the leaf-cutter ants. Leaf-cutter ants are a suitable testbed for reference-free size estimation, because their workers differ vastly in both size and shape; in principle, it is therefore possible to infer body mass—a proxy for size—from relative body proportions alone. Inspired by earlier work by E.O. Wilson, who trained himself to discern ant worker size from visual cues alone, we deployed deep learning techniques to achieve the same feat automatically, quickly, at scale, and from reference-free images: *Wilson Only Looks Once* (WOLO). Using 150,000 hand-annotated and 100,000 computer-generated images, a set of deep convolutional neural networks were trained to estimate the body mass of ant workers from image cutouts. The best-performing WOLO networks achieved errors as low as 11 **%** on unseen data, approximately matching or exceeding human performance, measured for a small group of both experts and non-experts, but were about 1000 times faster. Further refinement may thus enable accurate, high throughput, and non-intrusive body mass estimation in behavioural work, and so eventually contribute to a more nuanced and comprehensive understanding of the rules that underpin the complex division of labour that characterises polymorphic insect societies.

## 1 Introduction

Image-based size estimation is an important computer vision task, rendered challenging by the complexity and variability of visual cues. Applications range from agriculture and robotics to animal behavioural research [1–8]. Although different in motivation, these applications have in common the need to unify the appearance of the target subjects across images through tightly controlled imaging conditions, and to provide preprocessed information for accurate inference [1, 3, 7, 9]. Popular methods include convolutional network architectures that produce intermediate pose estimates, or binary image segmentations; both provide approximate measurements from which size estimates can be extracted [5, 6, 8]. In agricultural settings, e.g. in fruit processing plants [2, 4] or in livestock rearing [5, 6, 8], the recording environments are typically standardised, so that images have consistent camera-subject angles and camera-subject distances, which reduces task complexity. However, visual information by itself is still often insufficient to accurately estimate size, and it is therefore common practise to also include absolute scale information [1, 3, 7, 9], e. g. in form of reference objects of known size. Radar, sonar, or infrared light can help address the same problem, because mass can then be estimated from coarse 3D object reconstructions [10–12]. However, completely reference-free size estimation is rare [6].

The key challenge inherent in reference-free size estimation is that visually similar objects may well be of vastly different sizes; a tiny toy car can be readily confused with a real-sized car, through manipulation of image magnification and perspective [3]. As a consequence, global cues are usually vital for robust size estimation, but they cannot always be provided and, at the very least, limit application versatility. One scenario where reference-free size estimation should—at least in principle—be possible is where object size co-varies with object shape; size may then be inferred solely from the object itself through assessment of relative subject proportions. In this work, we tackle one such example: the workers of eusocial leaf-cutter ant colonies [13–16].

Leaf-cutter ants (Tribe Attini, Smith, 1858) form complex societies comprising large numbers of sterile “worker” ants that can vary by more than two orders of magnitude in body mass [13–15, 17–19]. Leaf-cutter ant colonies present a textbook example of a division of labour that transcends the ancient split into reproductive and sterile castes: morphological differentiation within the sterile individuals is coupled with task specialisation [14, 20]. The smallest individuals (minims) primarily tend to the fungus garden, the queen, and the brood; medium to large-sized workers (medias) cut and process plant matter; and the very largest workers (majors, often referred to as soldiers) almost exclusively engage in colony defence [14, 15, 18, 21]. Additional complexity arises within *Atta* colonies, as the variation in worker size is continuous [14, 22–24], and because the tasks carried out by workers of different sizes may change with the colony feeding state, age, distance to food sources, temperature, and ontogeny of individual workers [14, 17, 23, 25–28]. The materialised task preferences are hypothesised to lead to an ergonomic optimum—that is, workers are allocated such that each task is carried out to maximise the energy available to the colony [15, 29–32]. To give but a few examples, size frequency distributions of foraging parties appear to be adapted to the specific requirements of the available food sources [14, 33], and are affected by food source structural and mechanical properties [14, 17, 22, 34, 35].

Unravelling the “rules” that underlie the complex organisation of leaf-cutter ant colonies has been a long-standing objective in sociobiology, rendered challenging by the large number of involved behaviours and individuals per colony. In the absence of better options, researchers resorted to manual extraction and weighing of individual workers, which is time-consuming, error-prone, and disruptive (see, e.g. [13, 14, 36]). To minimise disruption, E.O. Wilson instead trained himself to estimate leaf-cutter head width by eye, aided by a physical lookup table in form of pinned workers [14]. Wilson reported that he was able to assign ant workers into one of 24 discretised size classes with an accuracy of 90 %, with the remaining ten percent placed into adjacent classes. This skill enabled Wilson to perform some classic experiments on the rules that govern division of labour in the leaf-cutter ants[14, 15, 18, 22, 37].

The goal of this work is to investigate to what extent a computer can be taught what Wilson taught himself, but without rich contextual information and from relative scaling properties alone. In making this attempt, we generate large and diverse training and validation datasets to enable future benchmarking and further development.

## 2 Methods

Our aim is to deploy deep learning-based computer vision approaches to automatically estimate the body mass of leaf-cutter ant workers from image cutouts without any absolute reference. To achieve this aim, training and benchmark datasets were curated, inference approaches implemented, and their performance evaluated; a small study with human participants was conducted to provide an indication of baseline performance. These steps are described in detail below.

### 2.1 Data collection and curation

A brief summary of the dataset collection and curation follows; a more detailed description can be found in the supplementary material (see 7.1).

#### 2.1.1 Training and Benchmark Datasets

Three datasets were collected (Fig. 1): (1) **MultiCamAnts**, a multi-animal dataset comprising images of ants on varied backgrounds, recorded with three synchronized cameras (Nikon D850, OAK-D, and Logitech C920), each with different perspective (Fig. 1 A-C); (2) **Test-A**, a single-animal dataset comprising images of individual ants on a neutral background, recorded with two synchronized cameras (Nikon D7000 and Logitech C920), each with different perspective (Fig. 1 D-F); (3) **Test-B**, a multi-animal dataset comprising images of a crowded foraging trail, representative of laboratory foraging experiments, recorded with a Luxonis OAK-D camera in top-down view (Fig. 1 G-I).

**Fig. 1.**
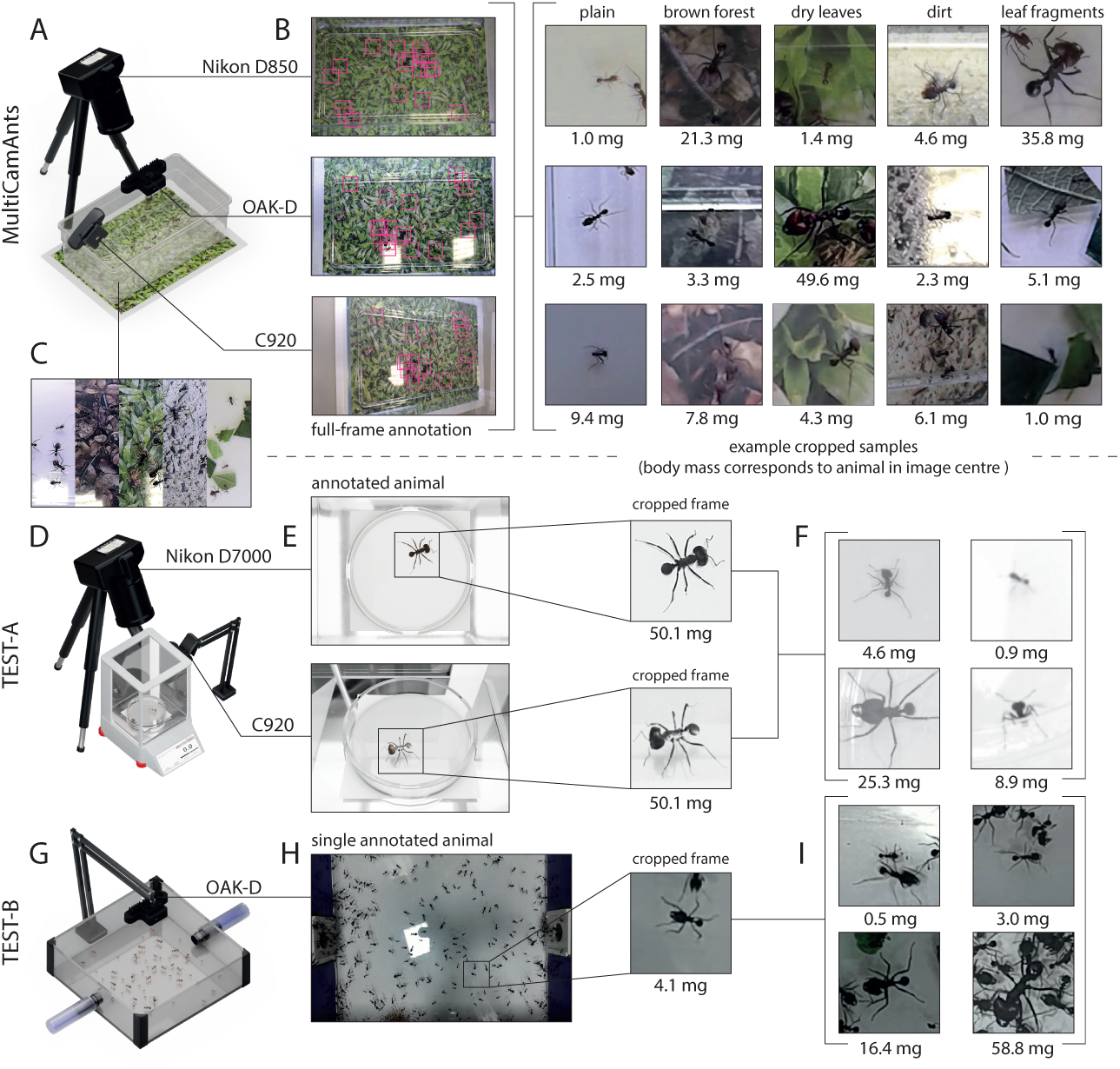
Collection and curation of training, validation, and out-of-distribution (OOD) test datasets. (A) Three synchronised cameras—a Nikon D850 with a 18-105 mm Nikkor lens, an OAK-D machine vision camera, and a Logitech C920—were used to record images of 20 *Atta vollenweideri* (Forel, 1893) leaf-cutter ant workers at 30 frames per second (fps). The 20 individuals were selected to represent 20 body mass classes, spaced equally across the mass range 1-50 mg in log-space. (B) The cameras recorded images from three unique perspectives, and with different magnification (Fig. 2, Supplementary Table 3 for exact worker masses, also available via Zenodo https://zenodo.org/records/11262427). 10,000 frames per camera were annotated semi-automatically for each of five recording scenarios, using *OmniTrax* [38]. (C) The recording scenarios differed in their image background, which was interchangeable to permit further variation: a default plain background, a textured brown forest floor, dry leaves, dirt, or a plain background with cluttered leaf-fragments. The 20 individuals always covered the same mass range, and represented the same 20 classes, but each background had a unique set of individuals. (B) The resulting dataset contained 3 *×* 5 *×* 10,000 = 150,000 labelled frames, each containing 20 individuals, resulting in a total of 3,000,000 cropped image samples (examples on the right). The first 80% of the full frame and cropped datasets were used as training data; the remaining 20% served as unseen validation data. Two out-of-distribution datasets were curated for further benchmarking. (D) Dataset *Test A* was recorded with a Nikon D7000 camera, equipped with a micro Nikkor 105 mm lens, oriented top-down, and a Logitech C920 camera at an angle of approximately 30% to the vertical. A single leaf-cutter ant worker was put into a Petri dish placed on an ultra fine scale, and (E) between 20 to 50 monochromatic frames were captured for each individual, resulting in (F) 4,944 cropped and annotated samples of 134 individual ant workers. (G) *Test-B* was recorded with an OAK-D camera positioned above a crowded container that served as a section of a laboratory foraging trail. (H) Individual workers were annotated with the manual tracking module of *OmniTrax* [38]; (I) a total of 30,526 cropped RGB samples of 154 individuals were extracted.

The **MultiCamAnts** dataset comprises five sets of 20 ants, collected from the foraging arena of a mature laboratory colony of *Atta vollenweideri* (Forel 1893) leaf-cutter ants to represent 20 body mass classes that cover the worker size range of 1–50 mg; class centre-to-centre distances were spaced equally within this range in log10-space to achieve a more fine grained class resolution among more common smaller worker sizes (Fig. 2 A-C). The visual appearance of the frames was varied by exchanging the arena background, and by scattering leaf-fragments, such as to emulate the appearance of foraging trails (see Fig 1 C). 150,000 labelled frames were annotated with both per-individual mass and bounding box data, exported as cropped samples, and split into 2.5 million training and 0.5 million validation samples (80/20).

**Fig. 2.**
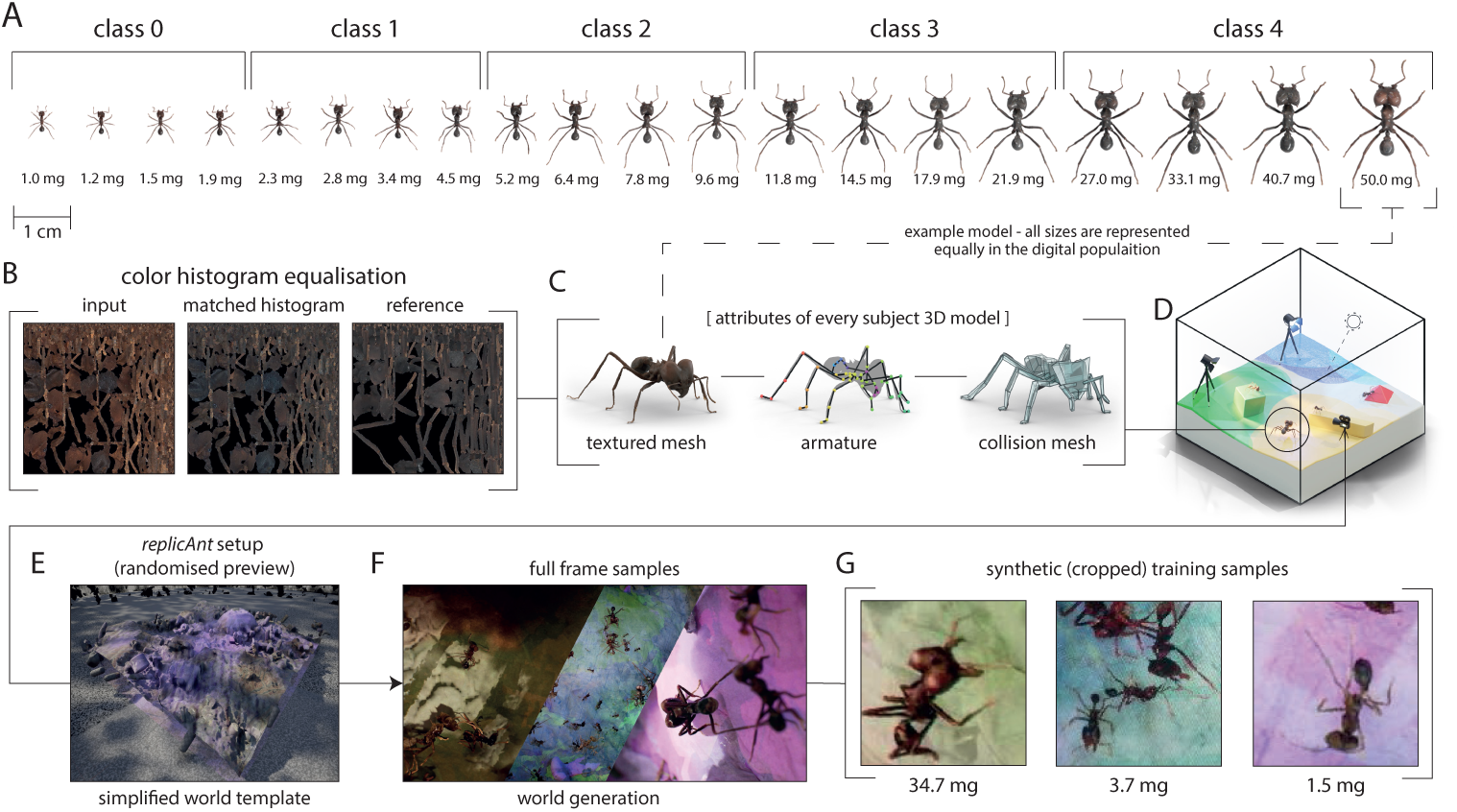
In an effort to increase prediction generalisability, real datasets were augmented with synthetic data, generated with *replicAnt*, an open source platform that places 3D models of animals in procedurally generated environments, and then produces annotated images from variable perspectives [39]. (A) “Digital twins” representing each of 20 leaf-cutter ant size classes were created with *scAnt* [40], an open source photogrammetry platform; for 5-classification, four individuals were grouped to form one class. (B) To remove colour variation caused by subtle differences in imaging conditions, colour-histogram equalisation was performed for all texture maps, using a custom-script written in python. (C) All 3D models were then retopologised and rigged in Blender (version 3.2), prior to porting them to (D) *replicAnt*, where a low polygonal collision mesh was computed, and the sample randomisation procedure configured. A digital population, comprising 200 individuals that differed in scale, hue, contrast, and saturation, formed the basis for (E) a large synthetically generated dataset, comprising (F) 100,000 full-frame examples. (G) Cropped training samples were extracted from the original full-frame samples using a data parser.

The **Test-A** dataset contains 4,944 cropped image samples of 131 ants, placed individually in a Petri dish on an ultra fine scale with a white background. The ants ranged in body mass from 0.5 to 25 mg, and were chosen at random from the colony foraging box (Fig. 2 D-F) Pictures were captured from two cameras (Nikon D7000 with a 105 mm micro Nikkor lens, Logitech C920) every 2 seconds.

The **Test-B** contains cropped samples of ants moving along a busy laboratory foraging trail, captured using a Luxonis OAK-D camera. 154 individuals were manually weighed and semi-automatically tracked using *OmniTrax* [38] to extract a total of 30,526 cropped image samples (Fig. 2 G-I).

#### 2.1.2 Augmentation with synthetic data

To augment the training split of **MultiCamAnts**, a synthetic dataset was produced using *replicAnt* [39], a multi-animal synthetic data generation pipeline implemented in Unreal Engine 5 (Fig. 2). The dataset was based on 20 3D models of leafcutter ants, again representing the 20 size classes. The models were created from dried and pinned workers using *scAnt* [40], an open photogrammetry platform, and subsequently rigged using Blender (version 3.2) to enable *in silico* pose variation. To account for slight differences in scanning settings between different worker sizes, image textures were unified in appearance using colour-histogram equalisation. 100,000 annotated full-frame images were generated, varying the digital environment, the model texture, and the combination of size classes present in the image; a total of 910,000 cropped-frame samples were automatically extracted from the full-frame data.

All cropped-frame training and benchmark datasets are archived on Zenodo (https://zenodo.org/records/11167521); full-frame datasets are prohibitively large, but can be obtained from the corresponding author. All custom tools and scripts used in this study are open source, and accessible on GitHub (https://github.com/FabianPlum/WOLO and https://github.com/evo-biomech/replicAnt).

### 2.2 Inference approaches

Two inference approaches were tested: image patch regression and classification. Regression is arguably the most natural implementation of the mass-estimation problem, but suffered from prediction bias and lower categorical accuracy (see results). Classification appeared less prone to such bias, at the cost of a unavoidable minimal error defined by the difference between ground truth sample vs class-centre mass.

#### 2.2.1 Regression

Regression was performed by a simple VGG-style[41] convolutional neural network, consisting of three blocks with increasing depth (32, 64, 128 filters respectively); dropout layers and batch normalisation were added as regularisation elements (See Supplementary Table 1). A single output node was followed by a sigmoid scaling layer, to restrict the range to 0 to 1, and subsequently remapped to the original mass range to extract the network’s predictions (see Supplementary Table 1).

Regression was conducted on 128 128 3 image samples (resolution in x, resolution in y, colour channels), cropped such that the thorax of the target animal was located in the image centre (Fig. 3 A). Networks were trained to minimise the absolute prediction error, i. e. the loss function was the Mean Squared Error (MSE):

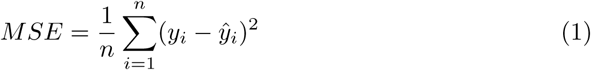

**Fig. 3.**
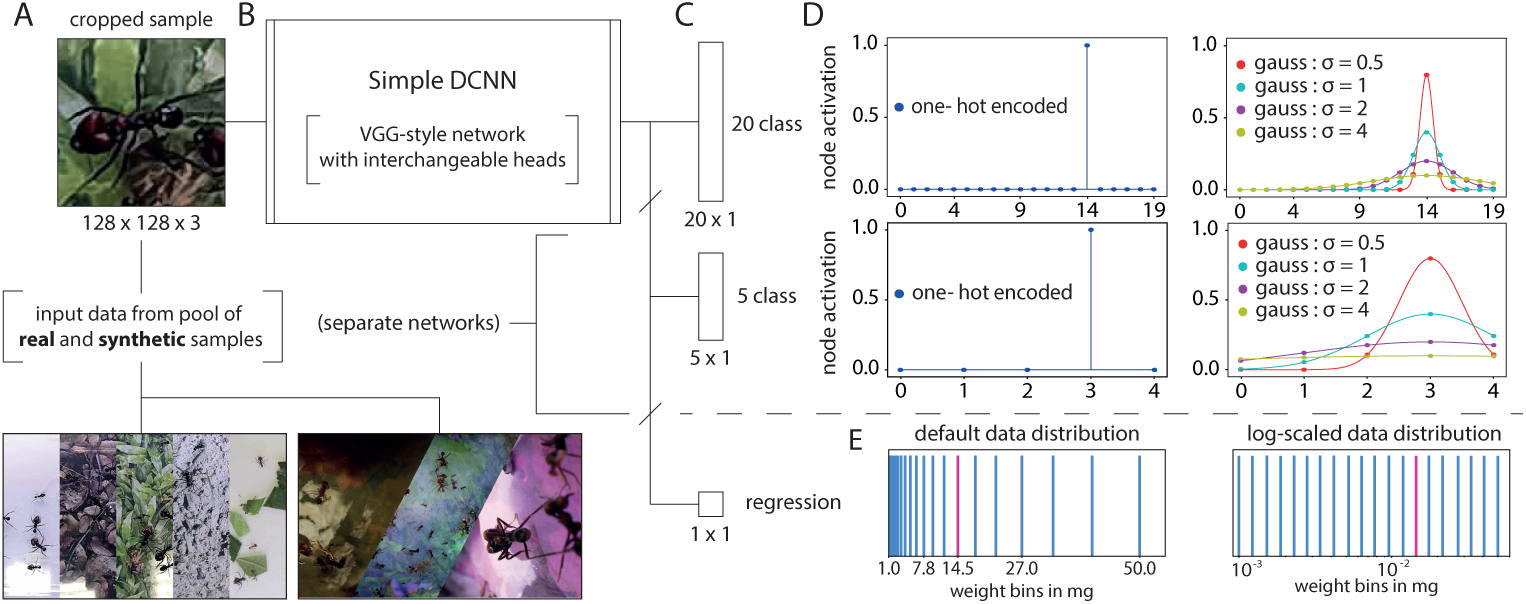
Schematic overview of deep convolutional neural network (DCNN) architectures and training paradigms for the estimation of ant worker body mass from reference-free images. The figure depicts cropped-frame classification and regression (A) 128 *×* 128 pixel cropped-frame samples were extracted from annotated images, and fed into (B) a VGG-style network [41] with (C) interchangeable heads to permit classification and regression. Networks were trained from scratch, using categorical cross-entropy loss for the classifier, and Mean Squared Error (MSE) loss for the regressor (see Supplementary Table S2 for details, available via Zenodo https://zenodo.org/records/ 11262427). (D) The relationships between classes at training time were encoded by assigning either default one-hot encoded, or custom class-aware Gaussian label smoothing to the classifier’s output layer. Label smoothing was implemented to introduce a differential penalty for misclassification as a function of the distance between ground truth and assigned class; assigning a class 1 worker into class 5 then carries a bigger error than assigning it into class 2. (E) Output activations for the regressor were normalised at training time, and training was conducted both on absolute and log10-transformed labels to minimise the absolute or relative error, respectively.

Here, *n* is the sample number, *y_i_*is the ground truth, and *y*^*_i_* the prediction. A reasonable alternative is to minimise the relative error, realised by training networks on log10-transformed body masses (Fig. 3 D).

Networks were implemented in Tensorflow (v2.9.1), trained for 200 epochs using Adam optimiser [42] with a learning rate of 0.0001, informed by preliminary trials that identified the approximate onset of overfitting. In each epoch, the network saw each sample once, with a batch size of 1024, leading to approximately 2500 or 3500 parameter updates per epoch, for networks trained on real data and on mixed datasets, respectively.

#### 2.2.2 Classification

Classification was performed with the same network architecture as regression, with the sole difference that the sigmoidal scaling layer was replaced by a classifier head with SoftMax activation. Two classifiers were trained; one with 20 and one with 5 classes. 20 classes roughly match the discretisation used by Wilson [14], and 5 classes provided a suitable reference for comparison to human performance on reference-free images (see below). In both cases, classes span the mass range [1, 50] mg, and class centres were chosen such that class-centre masses were approximately equidistant in log10-space. Both classifiers were implemented in Tensorflow (v2.9.1), and trained for 50 epochs with a batch size of 1024, using the Adam optimiser [42], a learning rate of 0.0001, and categorical cross-entropy as the loss function.

A key difference between classification and regression is that all classification errors are equal: miss-classifying a 1 mg ant as a 100 mg ant results in the same classification error as miss-classifying it as a 2 mg ant; biologically, these errors are however very different. To distinguish between classification errors, we implemented a simple class relationship-aware Gaussian label smoothing algorithm. Unlike default one-hot encoding, the label smoothing method lifts the activation of adjacent classes to the target class *µ*, according to a normalised Gaussian distribution with standard deviation *σ* (see 3 D). The normalised activation *y*(*x*) of each output node was defined as:

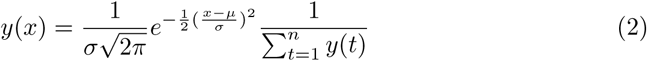

Label smoothing thus penalises incorrect predictions into a target class with a class-centre mass close to the ground truth less than an incorrect prediction into a class far away from it; it so renders categorical classification more similar to ordinal regression. Classifiers were trained with either default one-hot encoded labels, or class relationship-aware Gaussian label smoothing with *σ* = (0.5, 1, 2, 4).

### 2.3 Evaluation

We evaluated inference prediction error, categorical accuracy, and prediction stability. Accurate networks with low prediction errors predict body masses close to the ground truth (regressors), or the correct size class (classifiers); networks with high prediction stability predict the same absolute body mass or class for the same individual across frames.

#### 2.3.1 Prediction error and accuracy

Prediction error was assessed on the predictions averaged across all frames of the same individual. Regressors output continuous variables, and their prediction error was thus assessed on arithmetic means; classifiers, in turn, return categorical variables, and their prediction error was thus assessed on modes.

Prediction error was assessed via the Mean Absolute Percentage Error, or, in short, the prediction error (MAPE; also sometimes referred to as Mean Absolute Percentage Deviation (MAPD)):

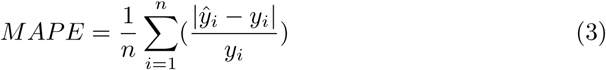

Here, *n* is the sample size, *y_i_*is the ground truth mass, and *y*^*_i_* is the estimated mass. A perfect regressor scores a MAPE of 0, and misclassifying a 1 mg as a 1.5 mg worker results in a prediction error of 50%—the same as misclassifying a 10 mg as a 15 mg worker. However, classifying a 1 mg worker as a 10 mg worker is associated with a prediction error ten times larger than classifying a 10 worker as a 1 mg worker. From these examples thus emerges a caveat that requires comment: because the MAPE quantifies relative instead of absolute errors, it penalises asymmetrically. As a consequence, unless the networks achieve high prediction confidence, they may learn to favour the prediction of small over large body masses, so leading to prediction bias and potentially even model collapse [43–45]; this is the main reason that MAPE was not used as a loss function during training.

The natural classifier performance metric is not a MAPE, but the categorical classification accuracy: the ratio between the number of correct classifications divided by the total number of classifications. A key weakness of using categorical accuracy as a metric to assess prediction performance on continuous data is that all misclassifications are treated identically, regardless of the relative error they carry; accuracy is thus of less relevance for the main aim of this study, which demands low prediction errors—but not necessarily high accuracy. Nevertheless, to facilitate direct comparison between classifiers and regressors, the regressor output was translated into a 20-class classification output by assigning each prediction the closest equivalent class centre in linear space, and its accuracy computed.

In comparing regressor and classifier performance, one further aspect requires comment. Because the MAPE is defined with respect to the ground-truth value, but classification only returns class centres, classifiers carry an unavoidable error that arises from mass discretisation: a classifier with perfect accuracy does not achieve a MAPE of zero. Instead, it has a finite prediction error that depends on the distribution of ground truth masses within each class. For the validation MultiCamAnts dataset, this error, *MAPE_ideal_*, was 6.14% and 22.75% for 20-class and 5-class inference approaches, respectively.

#### 2.3.2 Prediction Stability

The prediction stability and precision of predictions for continuous variables is best assessed via the coefficient of variation (CoV)—the ratio between sample standard deviation and arithmetic mean. However, no CoV can be defined for classifiers, so rendering comparison between regressors and classifiers impossible. To allow comparison, we instead again transformed regressor output into the 20-class equivalent (see above), and then assessed precision for both regressors and classifiers as the ratio between the number of samples assigned to the prediction mean or mode respectively *ỹ_i_*, and the total number of predictions *m_i_* for the same individual *i* across all frames *j*:

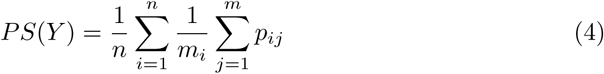

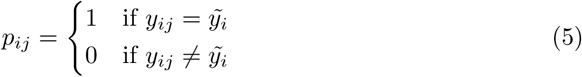

A network with high precision thus achieves a prediction stability of unity. Accuracy, prediction error, and prediction stability can also be assessed qualitatively via confusion matrices, provided for both class-equivalent regressor and classifier output.

### 2.4 Human performance

In order to provide an approximate performance baseline, 14 colleagues with and without experience in leaf-cutter ant research were asked to participate in an anonymised mass estimation study (Supplementary Table S3, available via Zenodo https://zenodo.org/records/14747391).

A simple online survey was implemented in SoSciSurvey [46]), to measure human performance on the 5-class cropped-frame mass estimation task (Fig. 8). Participants were first briefed on the purpose of the study and the upcoming task. They then agreed to the study conditions, and self-declared whether they work regularly with leaf-cutter ants. Next, participants were shown a simple task description. Akin to the way Wilson used a physical lookup table [14], participants were shown a digital lookup table as a guide (Fig. 2 and Fig. S 8 B-D). Every participant was initially shown 20 training examples in randomised order; after providing a classification, the correct size class was revealed. The 20 images were sampled from each size class of the original MuliCamAnts training split (see Fig 1 A-C), ensuring that participants saw one sample from each size class. The training phase was followed by a test phase during which randomly sampled cropped frames from all three datasets were shown—10 from each dataset, for a total of 30 test samples. At this stage, no further feedback was provided, and all classifications were recorded for later evaluation.

A full overview of all dataset combinations, network training strategies, and comprehensive performance evaluation on all validation and test data is provided in **Supplementary Table 3** (available also via Zenodo https://zenodo.org/records/14747391).

## 3 Results and Discussion

Estimating body mass from a single image is a challenging task, and usually requires the provision of reference lengths or other cues. In *Atta* leaf-cutter ants workers, body size variation is accompanied by changes in body shape [25, 47–53]; it thus ought to be possible to learn how to estimate body mass without external cues, and for images with variable magnification [14]. To achieve such reference-free mass estimation, we trained a variety of deep convolutional neural regressors and classifiers, and assessed their categorical accuracy, prediction error, and prediction stability. For the sake of clarity and brevity, we here only summarise the main trends and key results; a comprehensive overview is provided in Supplementary Table S2 (available via Zenodo https://zenodo.org/records/11262427).

### 3.1 Regressors achieve intermediate prediction errors, and can be strongly biased

A regressor network trained on absolute body masses achieved a prediction error of 18.44 %, with an accuracy below 50%. This accuracy, however, was strongly biased: the smallest size class had an accuracy almost four time lower than the largest size class (Fig. 4, B). This bias is likely the result of setting the absolute error as the loss function; it should consequently be resolvable by demanding minimisation of relative errors, i. e. by training regressors on log10-transformed data instead (Fig. 3 E). A network trained on log-transformed data achieved comparable error and accuracy, but prediction bias appeared strongly reduced, supporting this conjecture (Fig. 4 C & D). The 20 class-equivalent prediction stability slightly increased for the model with log-transformed class labels (one-hot encoded labels, PS = 0.516; log-transformed labels, PS = 0.528).

**Fig. 4.**
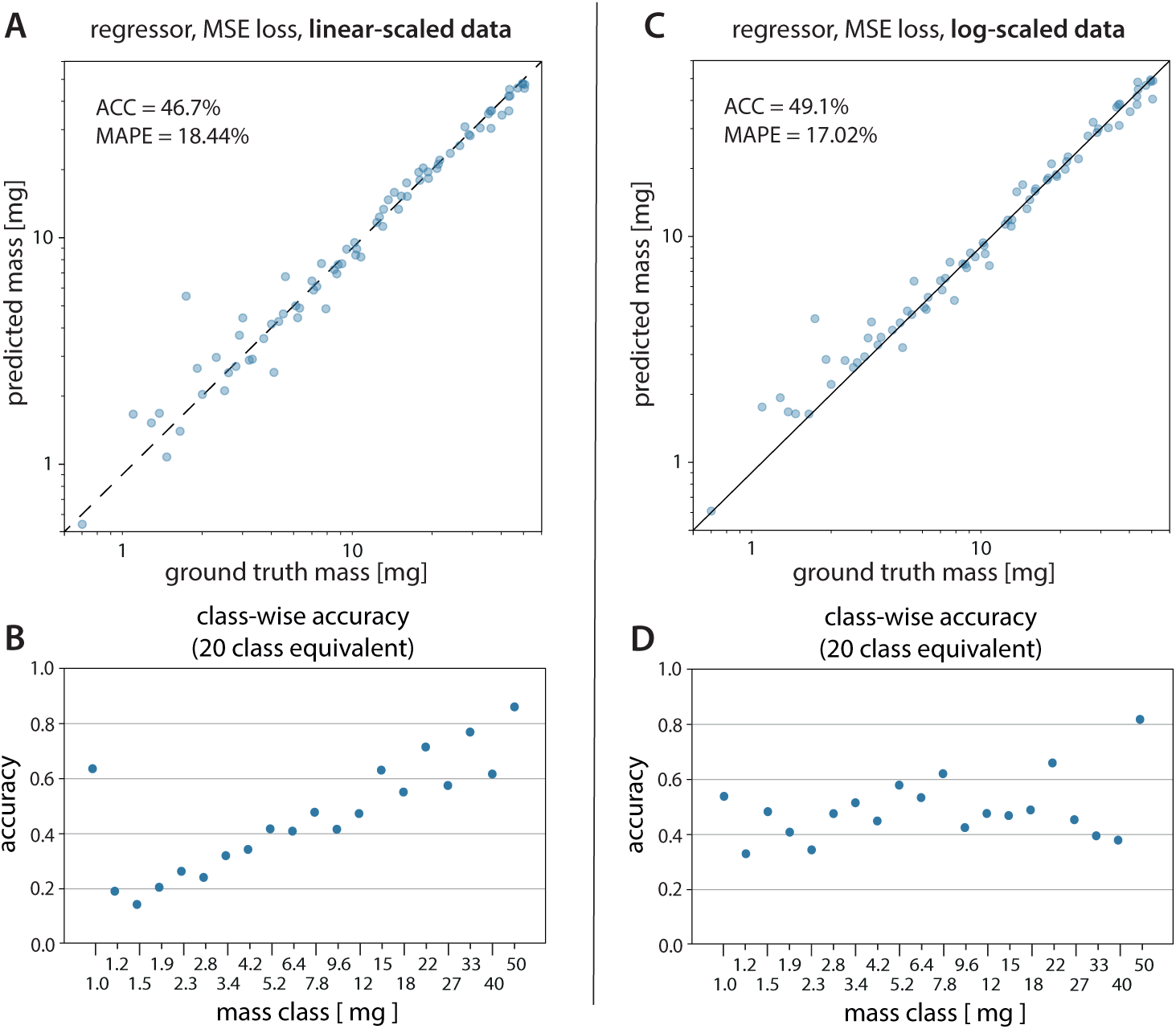
The most natural implementation of the mass estimation problem is a regressor network. (A) & (C) Deep regressor networks, trained on 2.5 million cropped frame samples and tested on 0.5 million samples from withheld within-domain data, achieved intermediate mean absolute percentage errors (MAPE) of about 20%, although predictions were generally clustered closely around the parity line (dashed) that indicates perfect prediction. Regressor networks trained on raw (untransformed) input data were also strongly biased, evident in a systematic variation of accuracy with size: mass predictions were almost four times more accurate for the largest compared to the smallest workers, a result most clearly visible when the regressor output is translated into a 20-class classifier equivalent, as shown in (B). (C) & (D) Accuracy bias was substantially weaker for regressor networks trained on log10-transformed body masses, i. e. when the loss function demanded minimisation of relative rather than absolute errors.

### 3.2 Classifiers achieve higher categorical accuracy, but suffer in prediction spread

Classifiers trained on one-hot encoded class labels achieved an error comparable to the best performing regressor, and, as expected, had a much improved accuracy: about 70% of all samples were assigned to the correct class (see also Supplementary Table 3). Classifiers also had noticeably higher prediction stability, when measured across estimates of the same individual. However, the price paid for these improvements was a reduction in prediction precision, as can be qualitatively assessed from the confusion matrices depicted in Fig. 5 B, D.

**Fig. 5.**
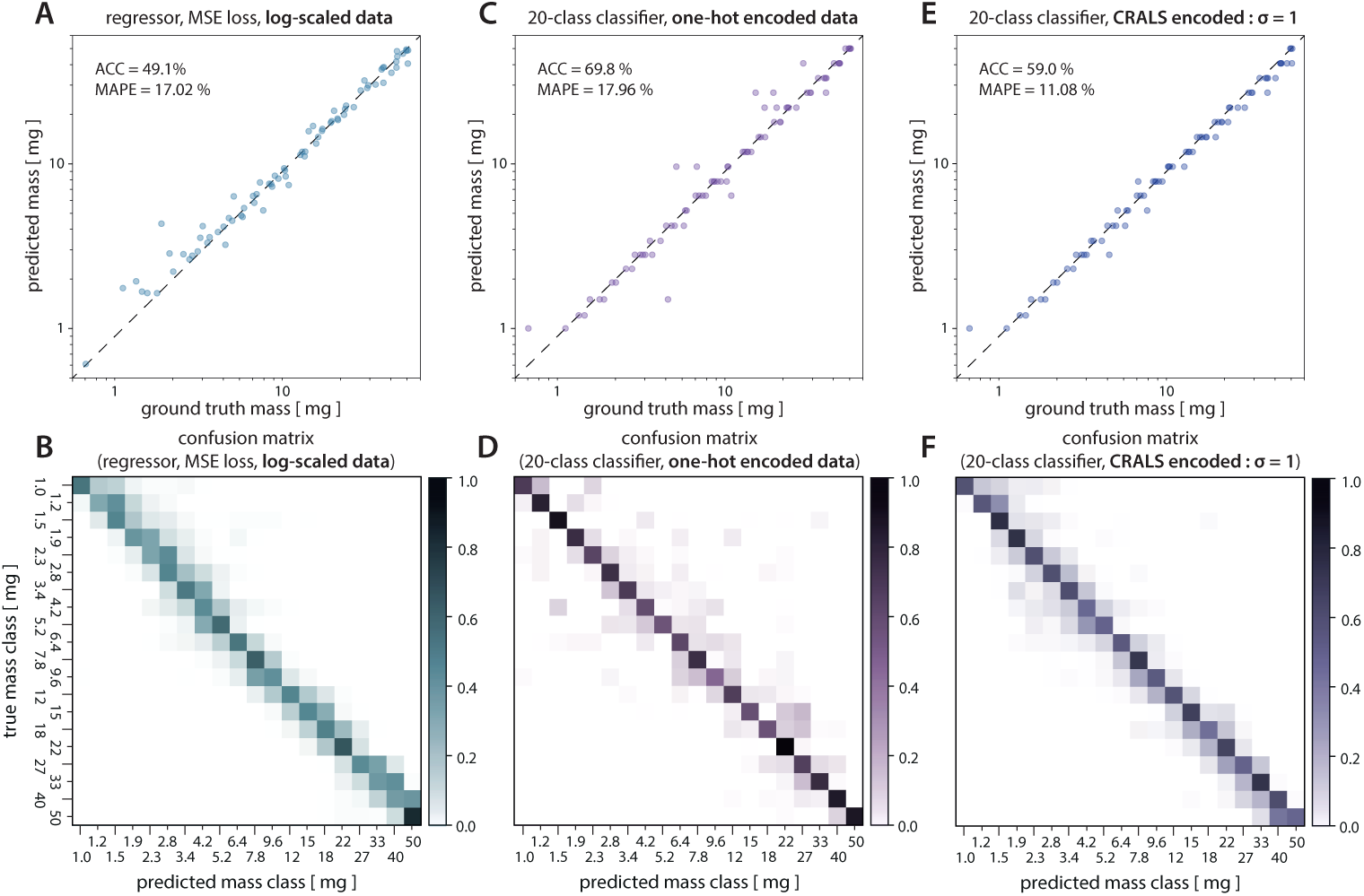
Classification presents an alternative method to size estimation. Classifiers achieved lower or comparable error, higher accuracy, had no obvious prediction bias, but were more prone to outliers. (A, C, E) Parity plots for within-distribution validation data of (A) the best regressor, trained on log10-transformed body mass data; (C) a classifier trained on 20 class discretised data with one-hot encoded labels; and (E) a classifier trained on the same data but with class-relationship-aware Gaussian label smoothing (*σ* = 1). Datapoints represent the arithmetic mean for the regressor, but the mode for the classifiers, evaluated across all frames for each unique individual. (B, D, F) Discretised confusion matrices corresponding to the parity plots. For the regressor, predictions clustered tightly along the parity line, but with a more pronounced spread around the class of highest activation. Classifiers, in turn, were more prone to outliers, but show practically no prediction bias. Class relationship-aware label smoothing (CRALS) smoothing can be introduced as a regularisation technique that controls the trade-off between these two effects, resulting in reasonably accurate, precise, and unbiased mass estimation.

### 3.3 Label smoothing combines the strengths of regressors and classifiers

Regressors generally achieved a lower accuracy, were prone to bias, but had good precision; 20 class Classifiers trained on one-hot encoded data achieved high accuracy and low bias, but suffered from lower precision. These differences likely reflect the reliance on ordinal vs categorial variables in regressors vs classifiers, suggesting a route to combine the best of both worlds: class-relationship-aware label smoothing (CRALS), effectively acting as a regularisation technique (see methods). The best network trained with CRALS achieved prediction errors as small as about 11%—close to the best-case prediction error of 6% that is associated with the discretisation of continuous data into 20 discrete classes (see methods). At the same time, accuracy suffered only slightly, and the number of outliers was reduced: predictions were clustered closely around the mode, across all size categories (cf. Fig. 5 B, D and F).

### 3.4 Performance on out-of-distribution data

The best network—a 20 class classifier with CRALS, *σ* = 1, achieved an error of about 11% on validation data—so low that it is likely sufficient to enable meaningful work on leaf-cutter ant foraging behaviour[15, 18, 53–56]. But any such network must ideally be able to generalise—that is retain performance on out-of-distribution (OOD) data without further refinement. On Test B, the same classifier had about four times the prediction error (MAPE = 43.62 %), and was much noisier (Fig. 6 B). Performance dropped even more on Out-Of-Distribution (OOD) Test A; the prediction error increased about seven-fold, to 73.86%, and, categorical accuracy dropped to as little as 0.07 % in Test A (Fig. 6 A) Clearly, prediction robustness and network generalisability remained somewhat wanting. The most likely origin of this result is overfitting, or to be precise, reliance on recording-specific cues that allow for high performance on unseen within-distribution data; other potential sources of error arise from different colour spaces in the recorded datasets, motion blur and 3D pose variation (TEST A), as well as a higher degree of individual overlap and occlusion (TEST B).

**Fig. 6.**
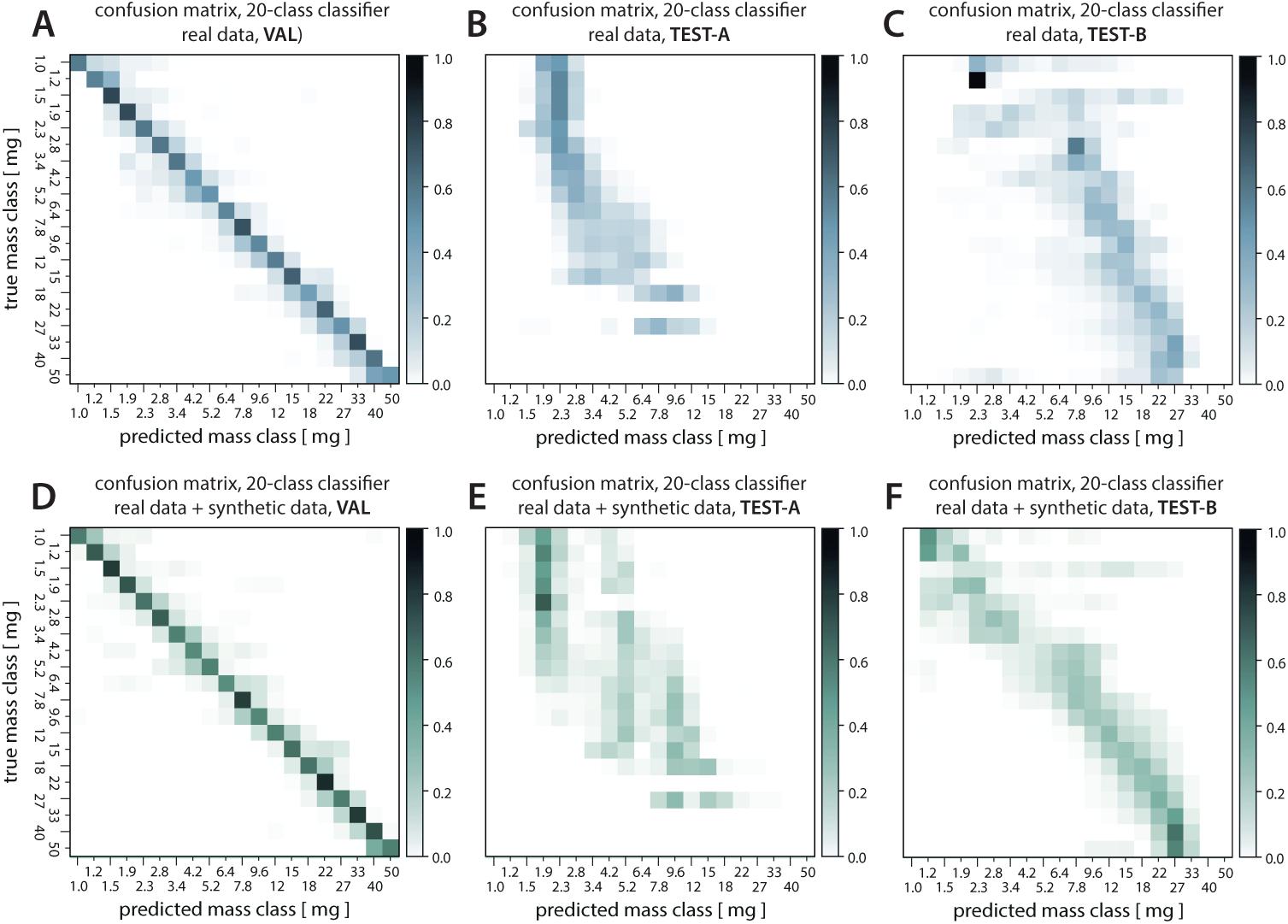
The ideal mass estimator generalises to unseen scenarios. However, even the best network— a 20-class classifier trained with Class-Relationship Aware Label Smoothing (CRALS)—showed a significant performance drop when put to work on out-of-distribution (OOD) data, as indicated by the parity plots (A, B). One way to address this problem is to augment the training data with diverse computer-generated synthetic data [57–60]. In support of this idea, synthetic data, generated with the dedicated open source tool replicAnt [39], helped networks retain their qualitative accuracy, and noticeably improved their quantitative accuracy. Data points represent the prediction mode per individual.

### 3.5 Synthetic data increase prediction robustness

To improve the network’s ability to generalise to out-of-distribution (OOD) data, the training dataset was supplemented with computer-generated synthetic images. This augmentation had only a small positive effect on validation performance, but strongly improved OOD performance (Fig. 6, and Supplementary Fig. S3): prediction errors dropped by about 20%, and categorical accuracy improved by about 5% for both Test A and Test B (see also Supplementary Table 3). Test A remained challenging, with even the best networks showing a prediction error as high as 55%, perhaps because its images provide next to no contextual information. For the same reason, they are, however, also somewhat artificial constructs; few, if any, real use cases will resemble these imaging conditions.

### 3.6 The best networks outperform humans

Human participants generally performed slightly worse than the best implemented networks in terms of prediction error and categorical accuracy; experts performed consistently better than non-experts (Fig. 7 A-C). Due to the small sample size, human bias cannot be reliably estimated. Wilson reported an accuracy of 90 % on a classification task with 24 classes, an accuracy not achieved by any of our participants or trained models, even in coarser classification tasks[14] ([see also 39]). It is unclear to what extent this difference in performance stems from sheer training, innate skill, from richer contextual information, or from a combination of all of these: Wilson observed foraging workers for prolonged periods, and had a detailed physical lookup table at hand. In contrast, human participants only received a short training, and a single low-resolution cropped image. This pilot study thus cannot provide a reasonable indication of the upper limit of human performance; but we believe that it supports the weaker conclusion that the task itself is hard, as well as the assertion that the trained networks learned a considerable amount from the training data provided.

**Fig. 7.**
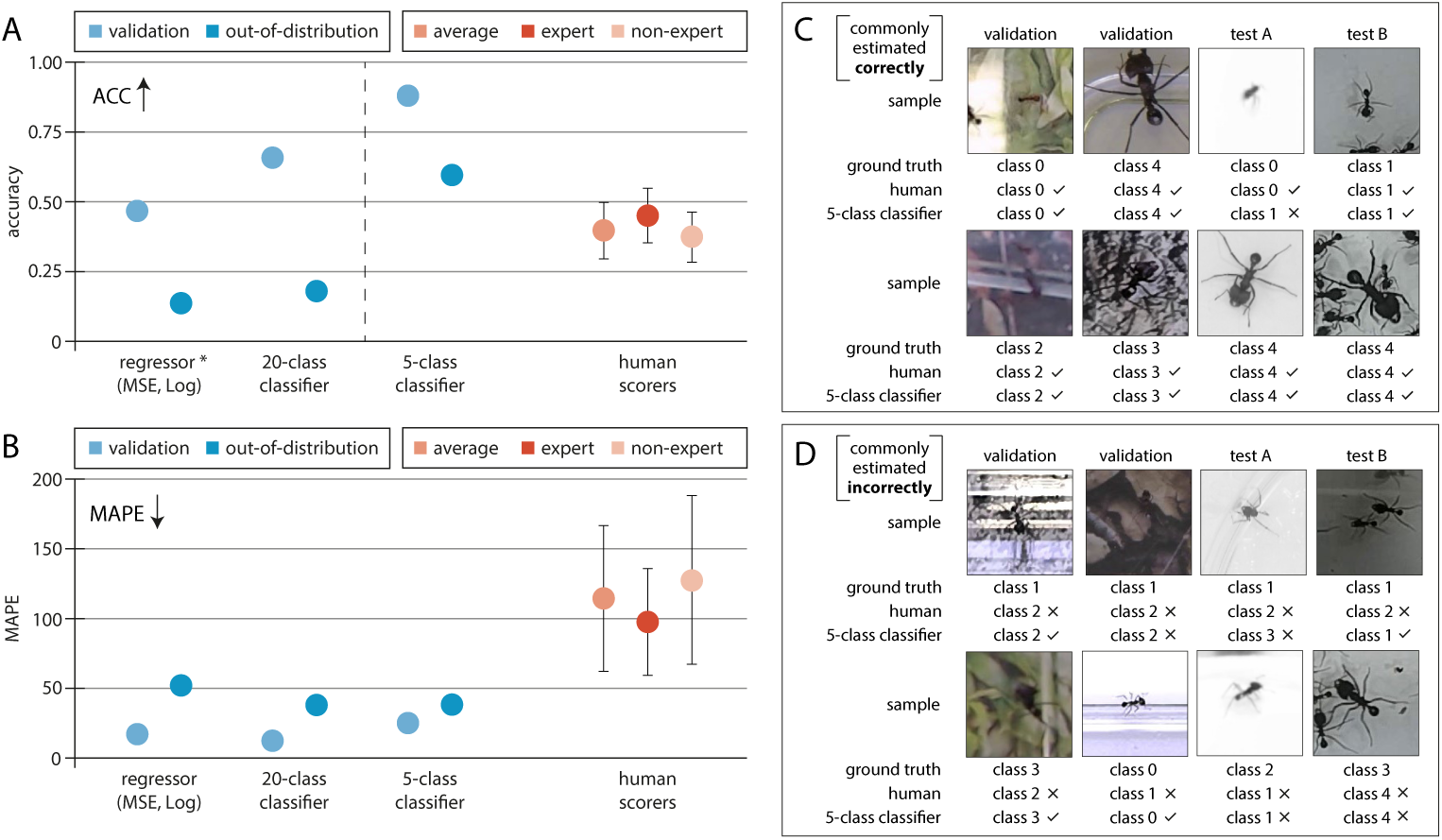
To place network performance into context, a small human predictor study was conducted. (A) The best networks achieved (A) similar or higher accuracy, and (B) lower prediction errors than humans.(^∗^Note, that humans performed a 5 class classification task, thus categorical accuracy of the regressor and 20 class classifier appear deflated by comparison, due to the finer class granularity.) Collections of representative image samples that were typically classified (A) correctly and (D) incorrectly by human annotators, compared with the classification of the best-performing 5-class classifier. Note that the computational models have seen substantially more training samples; combined with the small number of human participants (*n* = 14), this pilot study thus serves to provide rough estimate of human performance to highlight the difficulty of reference-free mass estimation.

In an effort to understand the limits to both human and network performance further, it is instructive to inspect image samples that were frequently classified correctly or incorrectly, respectively (Fig. 7 D-E). Human participants and network variants struggled with similar images: workers that were out-of-focus (Fig. 7 D, bottom row, third column), excessive image noise (Fig. 7 D, top row, first column), unusual poses, or deviations from top-down perspectives that distort or obscure morphological landmarks such as head widths or leg lengths (Fig. 7 D, top row, third column) all rendered the inference problem harder. A notable exception to this rule appear to be images of the very smallest ant workers (e.g. Fig. 7 C, top row, class 0), on which networks often performed poorly, but humans often did well, likely because they (correctly) inferred that the low magnification and high noise imply a small animal size. Although the same may be expected for networks, this is exactly one of the principal advantages of synthetic data, because it can avoid such bias, which otherwise may lead to overfitting.

Irrespective of prediction errors and accuracy, the main difference between human and computational classification lies elsewhere: Human annotators took on average six seconds to classify an image sample. The trained networks, in turn, performed about 1000 times as many predictions in the same time; they thus vastly outperform humans in terms of speed.

## 4 Conclusion and Outlook

Inspired by the work of E.O. Wilson, who trained himself to estimate leaf-cutter ant worker body size by eye [14], the aim of this work was to investigate the possibility of inferring ant body mass from reference-free images, using deep convolutional neural networks. Because size differences in leaf-cutter ants are associated with differences in shape, this aim ought to be achievable in principle.

Despite relatively large amounts of training data, the performance of even the best networks remained below the self-reported accuracy of E.O. Wilson; it was, however, comparable to that of human annotators that have had less dedicated and extensive training. The best networks had a size-independent (unbiased) prediction error of 11.94 % with an accuracy of 64.7 % across 20 mass classes, which may well be good enough for many research purposes[15, 18, 53–56]. The key advantage of the automated approaches is their vastly superior speed; any loss in accuracy can likely be balanced by an increase in sample size. Indeed, given that *Atta* trails readily contain thousands of individuals, automated mass-estimation may well be the only realistic and affordable option to obtain data on size-frequency distributions at the required scale.

Where mass estimation with even lower errors is needed, the most effective route may be the provision of an absolute scale, as done in previous related work [2, 3, 7]. Worker size can then be measured directly from the images, e. g., through body length or pixel number, with either pose estimators [61–63] or binary masks [2, 4, 5], respectively. However, for all its advantages, a direct measurement-from-pose approach is not without problems: parallax errors can only be managed through tight control of recording conditions, so reducing flexibility; and key body markers may often be occluded, be it by leaf-fragments carried or by other individuals that cross paths on busy foraging trails. Notwithstanding these difficulties, deep learning-based reference-free mass estimation has the potential to become a valuable asset in behavioural research on leaf-cutter ants and other polymorphic insects. Ample opportunity for algorithmic improvement exists, such as the inclusion of hierarchical information[64], or the use of intermediate layer activations of the network as feature extractors for ensemble learning strategies[65]. Nevertheless, the presented inference approaches present a promising step towards more nuanced, efficient, and non-intrusive methods in the study of the fascinating social organisation of complex insect societies.

Apart from its potential practical application in leaf-cutter ant research, our work yielded insights of relevance for reference-free mass estimation more broadly. First, class-aware label-smoothing as a regularisation technique can reduce error and improve prediction stability in ordinal classification tasks. Second, the addition of synthetic data can improve network robustness, i. e. teach networks to generalise better to unseen scenarios [see also 39]. Third, the choice of performance metric is not trivial, and each specific choice brings its own strengths and weaknesses. Future work will have to carefully and systematically address the problem of performance metric and model selection in effectively ordinal classification tasks.

## Supporting information

SUPPLEMENTARY TABLE 1 - All ant masses

SUPPLEMENTARY TABLE 1 - All network performances

SUPPLEMENTARY TABLE 1 - All human scores

## 5 Data availability

The code produced in this study is openly available on Github under an MIT License. All cropped frame datasets, 3D models, additional tables, and best performing networks are available on Zenodo under a Creative Commons Attribution 4.0 International License. The code repository includes detailed documentation and scripts required for reproducing the analyses, while the datasets contain all relevant input data used for model training and validation. Links to the repository and dataset can be found at:

- Code repository: https://github.com/FabianPlum/WOLO
- Datasets: https://zenodo.org/records/11167521
- 3D Models: https://zenodo.org/records/11167946
- Trained networks: https://zenodo.org/records/14746456
- Supplementary Tables: https://zenodo.org/records/14747391

For any additional inquiries, please contact the lead author.

## 6. Acknowledgements

This study was funded by the Imperial College’s President’s PhD Scholarship (to Fabian Plum) and is part of a project that has received funding from the European Research Council (ERC) under the European Union’s Horizon 2020 research and innovation programme (Grant agreement No. 851705, to David Labonte). The funders had no role in study design, data collection and analysis, decision to publish, or preparation of the manuscript. We thank Theo Humbeeck, who helped with the collection and annotation of the out-of-distribution dataset *Test A*.

## 7 Supplementary information

### 7.1 Datasets

#### 7.1.1 Training and benchmark datasets

Three datasets were curated (Fig. 1): (1) a complex multi-animal dataset, recorded with three different synchronised cameras, from three different perspectives, on five different backgrounds, and with varying degrees of leaf clutter (MultiCamAnts, Fig. 1 A-C); (2) a simpler single-animal test dataset, recorded with two different cameras, from two different perspectives, and on neutral background (Test-A, Fig. 1 D-F); and (3) a top-down multi-animal dataset, recorded with a machine-vision camera, oriented top-down above a busy ant foraging trail (Fig. 1 G-I). All cropped-frame training and benchmark datasets are available via Zenodo (https://zenodo.org/records/ 11167521). For full-frame datasets, please contact the corresponding author.

(**1**) The MultiCamAnts recordings served as the primary dataset. Three cameras—a Nikon D850 with a Nikkor 18-105 mm lens, a OAK-D machine vision camera, and a Logitech C920—were used to record images of ants that moved inside an acrylic container that served as recording arena (250 mm x 150 mm x 90 mm). Videos were time-synchronised by triggering a Nikon SB-700 AF Speedlight Flash Unit, once before animals were placed into the arena, and then again after 10,000 frames had been captured; these two time points were used to synchronise the videos using After effects (CC version 2023, Adobe Inc.). The visual appearance was varied by exchanging the arena background, and by scattering leaf-fragments, such as to emulate the appearance of foraging trails (see Fig 1 C).

Five sets of 20 ant workers were taken from the foraging containers of a mature laboratory colony of *Atta vollenweideri* (Forel 1893) leaf-cutter ants, housed in a climate chamber at 25*^◦^C* and 60% humidity; individuals were weighed with a precision scale (OBX-223 Ohaus Explorer Precision Balance, ± 0.1 mg resolution), and sampling continued until representative specimens for each of 20 body mass “classes” had been collected; class centres were chosen such that they were approximately equidistant in log10-space, and covered the mass range [1, 50] mg (see Fig. 2 B). 20 ant workers were placed in the recording arena for each background at a time, and in order of ascending body mass; subsequent identification was thus possible without the application of physical markers.

A total of 10,000 frames were captured for each of five recordings; one for each background, each with a different set of 20 individuals. These frames were subsequently annotated with *OmniTrax*, a deep learning-driven multi-animal tracking add-on for Blender [38]. Using user-guided semi-automatic tracking, all top-down recordings from the Oak-D camera were annotated. Subsequently, using a custom python script, the camera projections of the remaining views were solved, and the extracted homography was used to translate top-down tracks into the adjacent video perspectives. A total of 150,000 samples were labelled, exported and converted into the format required by the respective inference method (section 2.2). Mass estimation via detections demands full frames as input, which was facilitated with custom data parsers. Classification and regression, in turn, operate on cropped frames that contain only the focal individual. Cropped frames with customisable aspect ratio and resolution were exported from *OmniTrax* ; class and identity information were encoded in the filename. For full frame samples, a text file was generated for each frame containing the location, bounding box dimensions, and class of all visible animals. 3 *times* 5 × 10,000=150,000 samples for each of 20 individuals lead to a total of 3,000,000 cropped images. The first 80% (2,500,000) of these images were used as training-, and the final 20% (500,000) images as validation data. The splits were fixed to avoid inflation of the validation scores, as can occur when training and test sets contain time-adjacent frames that are visually similar.

(**2**) In order to curate dataset *Test-A*, a camera rig was built around an ultra-fine scale(OBX-223 Ohaus Explorer Precision Balance, ± 0.1 mg precision. Fig. 1 D-E). 131 ants were chosen at random from the colony feeding box, leading to a mass distribution that roughly resembles that of natural foraging parties, ranging from 0.5 to 25 mg. Individuals were placed, one at a time, into a Petri dish with a white background, centred on the scale. They were then filmed with a Nikon D7000 DSLR, equipped with a micro Nikkor 105 mm lens facing downward, and a Logitech C920, fastened to a custom-built mount and oriented with a 30° angle relative to the vertical (Fig. 1 D). Images were captured from both cameras, leveraging OpenCV [66] and libgphoto2; scale readings were recorded manually. Each camera captured 20 images per individual, with a low sampling frequency of 0.33 Hz, chosen to increase the postural variation across images of the same individual. The resulting dataset contained 4,944 cropped monochromatic image samples; about 300 images were discarded because individuals were entirely out of focus, or had unrulily escaped the recording setup.

(**3**) A *Test-B* dataset (see 1 G-H) was curated to obtain crowded images, resembling the conditions on a busy foraging trail. An OAK-D machine vision camera was positioned above a custom-built acrylic container (280 280 90 mm), connected in between the laboratory colony and a foraging box via a system of flexible PVC tubes (diameter 2 cm). Footage was recorded in multiple sessions over the course of four weeks, with a frame rate of 30 fps, and for a period of 20 minutes. During these recordings the colony was actively foraging on bramble leaves provided in the foraging box. Because leaf-cutter ant workers were allowed to enter and exit the container *ad libitum*, body masses needed to be determined by manual worker extraction and subsequent weighing with the Ohaus precision scale. One at a time, 154 individuals were weighed in this way, and semi-automatically tracked in-post for ~ 200 frames using *OmniTrax* [38]. A total of 30,526 cropped RGB frame patches were extracted using *OmniTrax*.

#### 7.1.2 Synthetic datasets

A large and diverse synthetic dataset, consisting of computer-generated images, was produced to augment the training split of MultiCamAnts, in an effort to increase network robustness on Out-Of-Distribution data [39]. Synthetic data were generated with *replicAnt*, a computational pipeline implemented in Unreal Engine 5 and Python [39]. *replicAnt* takes textured and rigged 3D models as input, and places simulated populations of these models into complex, procedurally generated environments. From these environments, computer-annotated images can be exported, which can then be used as training data for a variety of based computer vision applications, including classification, detection, tracking, 2D and 3D pose-estimation, and semantic segmentation.

To provide the required 3D input models, 20 worker ants, distinct from those used in the MultiCamAnts recordings but with comparable size, were sampled from the laboratory colony (Supplementary Table S4, also available via Zenodo https://zenodo.org/records/11262427). Specimens were sacrificed via freezing to produce “digital twins” with the open source photogrammetry platform *scAnt* [40]. Specimens were prepared such that they were biting down on either a needle or thin PLA filament, so that their mandibles did neither touch nor overlap; this facilitated digital positioning of the mandibles at a later stage (see below). Specimens were pinned in an upright position akin to their natural stance, and left to dry at room temperature for at least one week prior to scanning. This drying step ensured that the joints had sufficiently stiffened to prevent movement during scanning. Specimens were scanned with the *scAnt* hardware configuration described in Plum and Labonte 2021, the code version from the May 2023 (*dev* branch), and the default masking parameters of an updated stacking routine (https://github.com/PetteriAimonen/focus-stack). Specimens lighter than 4 mg were digitised using a 75 mm MPZ Computar lens, and a custom-built focus extension tube (see [40] for details); all other specimens were photographed using a 35 mm MPZ computar lens and a 5 mm C-mount extension ring. All models are available via Zenodo (https://zenodo.org/records/11167946).

All scans were performed with a colour-coded 5 5 5 mm cube in view to enable both colour calibration and rescaling of the resulting 3D models. Scans were photogrammetrically reconstructed with 3DF Zephyr lite (v2023.03), with photo-consistency meshing enabled to retain fine structural details. It is not trivial to quantify photogrammetric reconstruction accuracy. As an approximate guide, *scAnt* can resolve step-changes in height of about 100 µm with an error of around 10%; this error drops to less than 5% for steps of 500 µm [40].

Reconstructed textured meshes were exported as FBX (”Filmbox”) files, and subsequently imported into Blender 3.2, to complete basic mesh cleaning (see [40]), and to apply a standardised armature ([39]; Fig. 2 D). The rigged mesh was retopologised to decrease the number of vertices from *>* 100, 000 to 10, 000, substantially reducing the subsequent computational load. The rigged and retopologised models were then scaled to their original size, using the colour-coded cube as reference, and the appearance of image textures was unified using histogram equalisation (fig 2 C). All models were then brought into *replicAnt* using the *send2Unreal* plug-in.

Within *replicAnt*, a large synthetic dataset was produced from a simulated population of 200 individuals; 10 from each original model. Within each of the 20 size classes, a randomised scale variation of 10% was applied, so that adjacent mass classes did not overlap in absolute scale. To produce a simple dataset with high levels of texture variation, the default generation environment within *replicAnt* 1.0 was chosen, with 70% of the asset scatterers removed. 100,000 image samples were generated. *replicAnt* ’s multi-class YOLO parser was used to export full-frame samples, and a custom-written second parser produced 128 × 128 px cut-out samples for every animal in every synthetically generated frame. These cutouts were rescaled when animals occupied a larger area to ensure that the entire animal was visible, and the subject class was encoded in the filenames. Individuals that occupied small fractions of the cutout were centred, and basic up-sampling was applied such that the larger side of the bounding box corresponded to at least 10% of either the width or height of the cropped image. A simple conversion script automatically sorted samples into discrete size folders, using the class information provided in the file name, and so produced the file-structure required by *TensorFlow.dataset* (see below).

All custom tools and scripts used in the curation and generation of real and synthetic data are open source, and accessible on GitHub (https://github.com/FabianPlum/WOLO and https://github.com/evo-biomech/replicAnt).

### 7.2 Survey - Human mass estimation

**Fig. 8.**
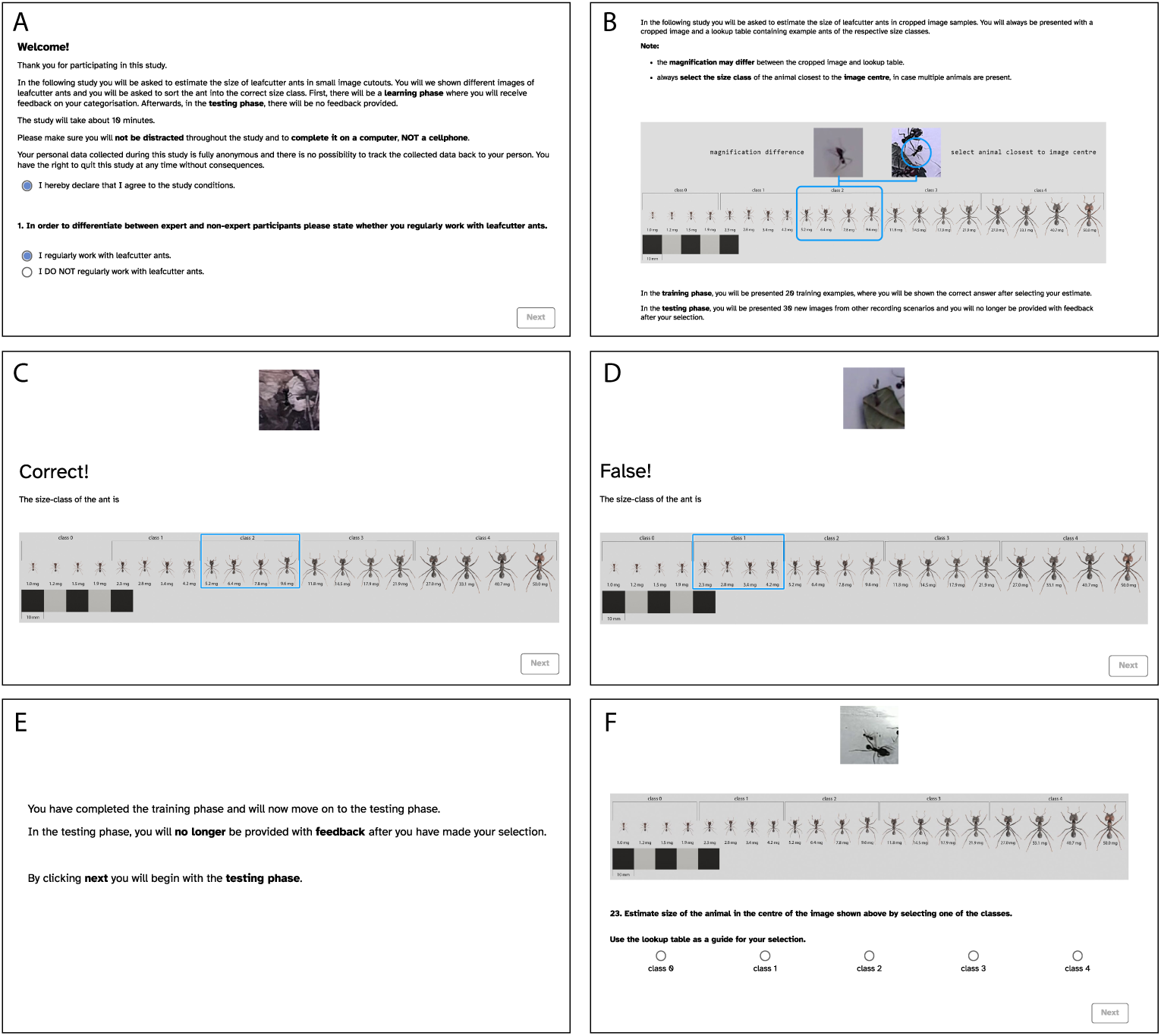
Main pages of the survey designed to measure human performance on 5-class mass estimation tasks. (A) Welcome page: participants are introduced to the study and prompted to agree to the study conditions, as well as to declare whether they work regularly with leaf-cutter ants (experts) or not (non-experts). (B) Task description: This page outlines the mass estimation task, and how participants are supposed to enter their answer; it also explains the differences between the training and testing phases of the survey. (C-D) Example pages from the training phase, shown after the participant has made a correct or incorrect selection, respectively. (E) This page is prompted after the participant has completed their training. (F) In the testing phase, no further feedback is provided after each estimate has been made.

### 7.3 Body mass as a function of body length

**Fig. 9.**
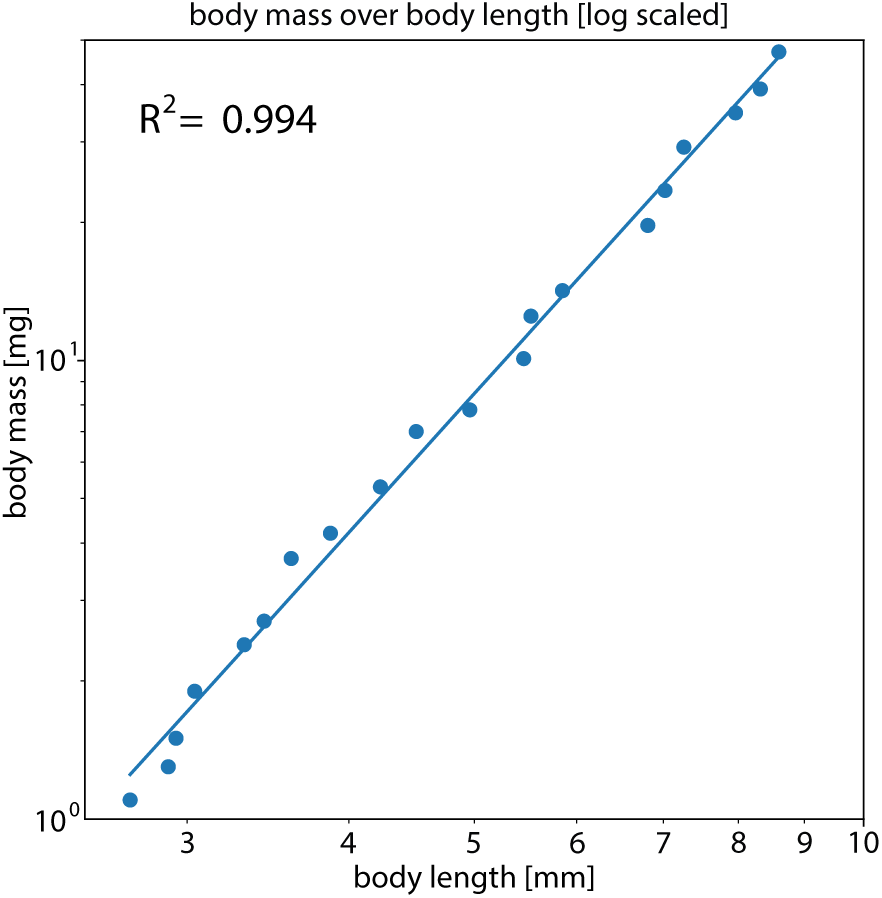
Body mass varies with body length [log scaled]. An ordinary least squares regression on log-transformed data yields an elevation of -1.25, and a scaling coefficient of 3.122, consistent with isometry.

### 7.4 Network architecture, reference-free cropped frame mass estimation

**Table 1.** CNN Architecture for cropped-frame regression and classification (Total parameters: 17,078,836 for 20 class networks)

| Layer (Type) | Output Shape | Parameters |
| --- | --- | --- |
| Input (input_layer) | (None, 128, 128, 3) | 0 |
| Conv2D (conv1.1) | (None, 128, 128, 32) | 896 |
| BatchNormalization (bn1.1) | (None, 128, 128, 32) | 128 |
| ReLU (relu1.1) | (None, 128, 128, 32) | 0 |
| Conv2D (conv1.2) | (None, 128, 128, 32) | 9,248 |
| BatchNormalization (bn1.2) | (None, 128, 128, 32) | 128 |
| ReLU (relu1.2) | (None, 128, 128, 32) | 0 |
| MaxPooling2D (pool1) | (None, 64, 64, 32) | 0 |
| Dropout (dropout1) | (None, 64, 64, 32) | 0 |
| Conv2D (conv2.1) | (None, 64, 64, 64) | 18,496 |
| BatchNormalization (bn2.1) | (None, 64, 64, 64) | 256 |
| ReLU (relu2.1) | (None, 64, 64, 64) | 0 |
| Conv2D (conv2.2) | (None, 64, 64, 64) | 36,928 |
| BatchNormalization (bn2.2) | (None, 64, 64, 64) | 256 |
| ReLU (relu2.2) | (None, 64, 64, 64) | 0 |
| MaxPooling2D (pool2) | (None, 32, 32, 64) | 0 |
| Dropout (dropout2) | (None, 32, 32, 64) | 0 |
| Conv2D (conv3.1) | (None, 32, 32, 128) | 73,856 |
| BatchNormalization (bn3.1) | (None, 32, 32, 128) | 512 |
| ReLU (relu3.1) | (None, 32, 32, 128) | 0 |
| Conv2D (conv3.2) | (None, 32, 32, 128) | 147,584 |
| BatchNormalization (bn3.2) | (None, 32, 32, 128) | 512 |
| ReLU (relu3.2) | (None, 32, 32, 128) | 0 |
| MaxPooling2D (pool3) | (None, 16, 16, 128) | 0 |
| Dropout (dropout3) | (None, 16, 16, 128) | 0 |
| Flatten | (None, 32768) | 0 |
| Dense (dense1) | (None, 512) | 16,777,728 |
| BatchNormalization (bn_dense1) | (None, 512) | 2,048 |
| ReLU (relu_dense1) | (None, 512) | 0 |
| Dropout (dropout_dense1) | (None, 512) | 0 |
| Dense (output) | (None, 20) | 10,260 |
| Softmax (classification) / Sigmoid (regression) | (None, num_classes) | 0 |

### 7.5 Gaussian label smoothing can reduce error in ordinal classification tasks

One hot-encoded labels are a sensible choice for categorical inference tasks. However, classes in discretised mass inference retain an ordinal characteristic, so that not all classification errors are equal. Consequently, penalising incorrect classification into adjacent classes less may help avoid overfitting, and increase the correlation between performance metrics in out-of-distribution data (see table 2). We trained various 5- and 20-class networks with a pre-trained Xception-net backbone (see https://github.com/FabianPlum/WOLO for information on which other backbones and training modes are supported). The resulting performance was lower than in our final presented work, which used a shallower network trained from scratch. The results shown here are therefore indicative of a trend in performance shifts with increasing levels of label-smoothing, not a performance optimum. The 5-class classifier, trained on mixed MultiCamAnts and synthetic data with class relationship-aware Gaussian label smoothing with *σ* = 2, achieved the highest overall absolute accuracy of 47.3% on out-of-distribution data. There also appeared to be a systematic decrease in the CoV, indicating increased classification precision. The best smoothing parameter *σ* varied with the number of classes; for 5 class models, the activation profile became flat, and performance in fact decreased for *σ >* 2 (see Table 2).

**Table 2.** Performance of 5 and 20 class classifiers trained with class relationship-aware Gaussian label smoothing, using an Xception-net backbone (NOTE: This is not the final network presented in the main paper and stems from preliminary trials). All networks were trained with mixed real and synthetic datasets. MAPE, categorical accuracy, Coefficient of Variation (CoV) are reported on validation data (500,000 unseen samples collected across the camera perspectives and background textures depicted in 1 (A-C)), and on out-of-distribution datasets A and B, comprising 4,944 and 30,526 samples respectively (see 1 (D-I).) Label smoothing increased both robustness and precision, as evident in increased performance metrics on out of distribution data, and a reduction of the CoV.

| classes | sigma | Validation scores |  |  |  | combined scores on Test A and B |  |  |  |
| --- | --- | --- | --- | --- | --- | --- | --- | --- | --- |
| | | SRCC ↑ | MAPE ↓ | acc. ↑ | CoV ↓ | SRCC ↑ | MAPE ↓ | acc. ↑ | CoV ↓ |
| 5 | 0* | 0.840 | 73.3 | 0.610 | 0.718 | 0.732 | 67.7 | 0.401 | 0.560 |
|  | 0.5 | 0.872 | 54.3 | 0.622 | 0.714 | 0.774 | 61.3 | 0.410 | 0.565 |
|  | 1 | 0.923 | 42.2 | 0.595 | 0.632 | 0.738 | 55.2 | 0.424 | 0.553 |
|  | 2 | 0.903 | 53.0 | 0.544 | 0.515 | 0.626 | 60.5 | 0.436 | 0.480 |
|  | 4 | 0.851 | 78.6 | 0.453 | 0.436 | 0.444 | 70.8 | 0.415 | 0.413 |
| 20 | 0* | 0.888 | 41.9 | 0.420 | 0.784 | 0.686 | 56.0 | 0.105 | 0.622 |
|  | 0.5 | 0.885 | 39.5 | 0.413 | 0.776 | 0.590 | 54.0 | 0.133 | 0.545 |
|  | 1 | 0.861 | 56.6 | 0.383 | 0.717 | 0.714 | 60.3 | 0.115 | 0.608 |
|  | 2 | 0.872 | 37.8 | 0.319 | 0.747 | 0.653 | 73.2 | 0.109 | 0.533 |
|  | 4 | 0.915 | 36.4 | 0.241 | 0.619 | 0.831 | 57.2 | 0.126 | 0.477 |

\* using default one-hot encoding without label smoothing.

### 7.6 Lower size-classes disproportionately affect MAPE scores

Regardless of inference approach, loss function, and label transformation technique employed, lower size classes disproportionately affect the overall MAPE scores, as evident from inspection of the class-wise MAPE scores (see 7.6), which were typically between 3 to 10 times higher for the smallest classes, even for overall well-performing inference approaches.

In addition to the size-dependence inherent in the definition of the MAPE score (see methods), the presence of larger individuals occluding the target animal in the same cropped frame likely inflates the error further. If inference approaches are selected according to the lowest MAPE, then a method that systematically underestimates body mass will do better than a method that systematically overestimates it: the MAPE favours models that underpredict the target distribution because it assigns more mass to data points with smaller ground truth values in the denominator, making these points more influential [43, 44]. Various additions have been suggested to counteract this property such as dividing the absolute error by the average of the predicted and ground truth value instead of the ground truth alone [44] or by log-transformation of the MAPE [43], and may be explored in future work.

**Fig. 10.**
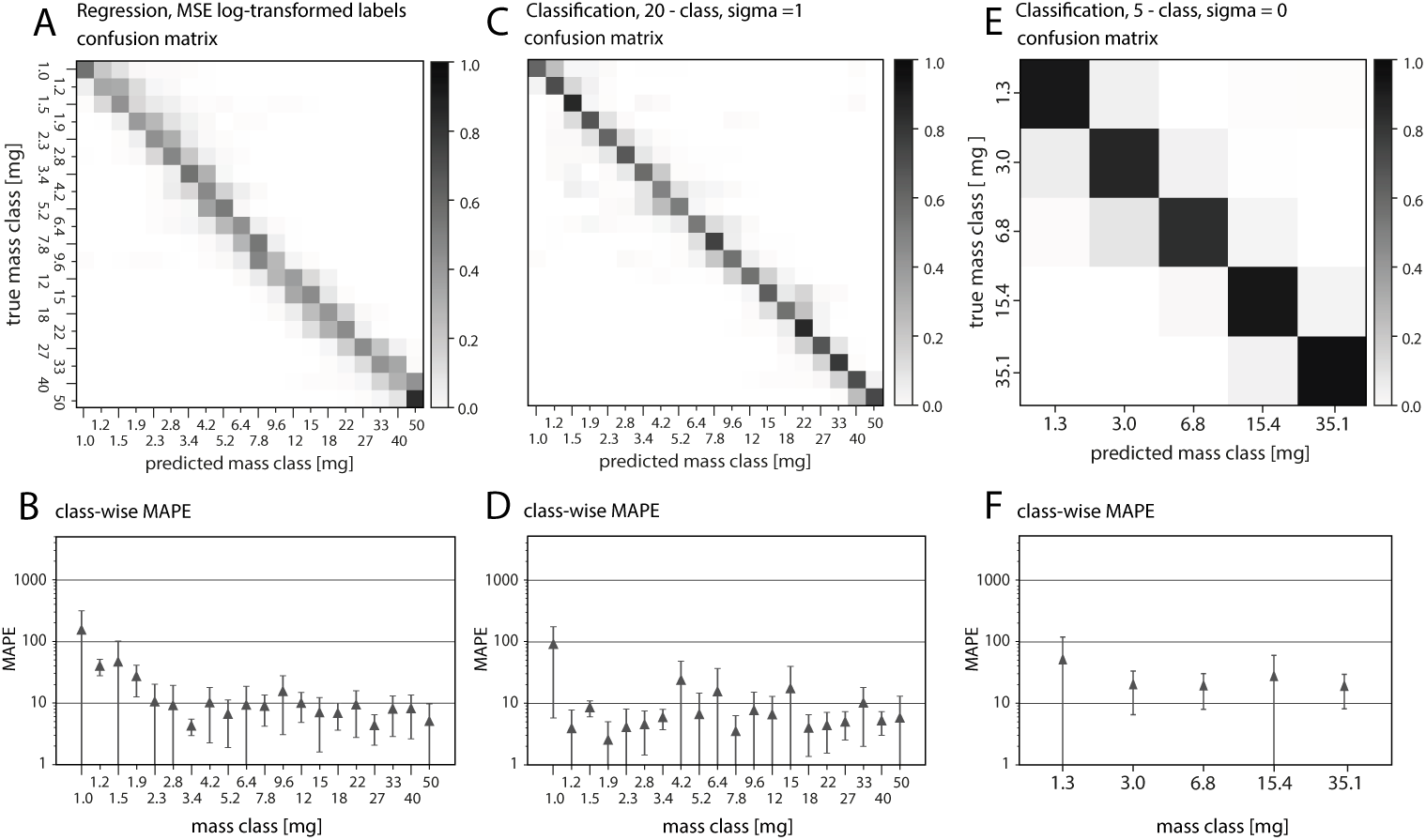
Mass estimation performance of classifiers and regressors. Accuracy on unseen validation data increases from left to right; all networks were trained on a mix of real and synthetic data. (A) A regressor, trained with log-transformed labels achieved a categorical accuracy of 0.460;

### 7.7 Results overview Table

**Table 3.** Performance overview of all reported models. . The accuracy, error (MAPE) and prediction stability are reported separately for predictions on unseen within-distribution (Validation) and two out of distribution test cases (Test A and Test B). For the regressors, accuracy is reported based on the 20-class-equivalent predictions.

| model name | Validation |  |  | Test B |  |  | Test A |  |  |
| --- | --- | --- | --- | --- | --- | --- | --- | --- | --- |
|  | ACC | MAPE | PS | ACC | MAPE | PS | ACC | MAPE | PS |
| **20 class classifier** |  |  |  |  |  |  |  |  |  |
| CL20_sigma-0_CE_VGG | 0.698 | 17.96 | 0.744 | 0.196 | 32.90 | 0.503 | 0.134 | 127.19 | 0.454 |
| CL20_sigma-0_CE_Xception | 0.393 | 42.62 | 0.459 | 0.064 | 51.40 | 0.239 | 0.101 | 69.02 | 0.524 |
| CL20_sigma-1_CE_VGG | 0.590 | 11.08 | 0.634 | 0.199 | 43.62 | 0.400 | 0.074 | 73.88 | 0.402 |
| CL20_sigma-1_CE_Xception | 0.270 | 34.04 | 0.367 | 0.084 | 50.91 | 0.172 | 0.074 | 69.00 | 0.372 |
| CL20_synth_sigma-0_CE_VGG | 0.700 | 13.16 | 0.758 | 0.161 | 39.40 | 0.546 | 0.132 | 71.09 | 0.412 |
| CL20_synth_sigma-0_CE_Xception | 0.414 | 42.62 | 0.476 | 0.076 | 60.96 | 0.205 | 0.104 | 61.91 | 0.368 |
| CL20_synth_sigma-1_CE_VGG | 0.647 | 11.94 | 0.679 | 0.245 | 23.00 | 0.402 | 0.115 | 54.41 | 0.345 |
| **5 class classifier** |  |  |  |  |  |  |  |  |  |
| CL5_sigma-0_CE_VGG | 0.864 | 24.74 | 0.881 | 0.543 | 55.27 | 0.702 | 0.546 | 41.21 | 0.688 |
| CL5_sigma-0_CE_Xception | 0.581 | 40.66 | 0.638 | 0.321 | 47.71 | 0.405 | 0.274 | 74.30 | 0.727 |
| CL5_sigma-1_CE_VGG | 0.853 | 23.54 | 0.877 | 0.487 | 52.86 | 0.722 | 0.434 | 82.89 | 0.722 |
| CL5_sigma-1_CE_Xception | 0.536 | 41.70 | 0.602 | 0.375 | 38.12 | 0.480 | 0.289 | 69.73 | 0.757 |
| CL5_synth_sigma-0_CE_VGG | 0.882 | 24.63 | 0.896 | 0.726 | 25.38 | 0.772 | 0.455 | 50.32 | 0.543 |
| CL5_synth_sigma-1_CE_VGG | 0.857 | 25.32 | 0.871 | 0.673 | 25.09 | 0.739 | 0.440 | 82.98 | 0.718 |
| **regressors** |  |  |  |  |  |  |  |  |  |
| REG_MSE_LOG_VGG | 0.491 | 17.02 | 0.528 | 0.202 | 38.81 | 0.362 | 0.138 | 68.73 | 0.183 |
| REG_MSE_VGG | 0.467 | 18.44 | 0.516 | 0.216 | 42.47 | 0.383 | 0.117 | 81.54 | 0.156 |
| REG_synth_MSE_LOG_VGG | 0.460 | 16.45 | 0.495 | 0.132 | 37.27 | 0.325 | 0.140 | 66.18 | 0.207 |
| REG_synth_MSE_LOG_Xception | 0.144 | 62.72 | 0.200 | 0.106 | 49.12 | 0.169 | 0.073 | 71.98 | 0.222 |
| REG_synth_MSE_VGG | 0.402 | 22.11 | 0.456 | 0.238 | 29.72 | 0.383 | 0.143 | 70.55 | 0.171 |

